# Sex-biased gene expression across tissues reveals unexpected differentiation in the gills of the threespine stickleback

**DOI:** 10.1101/2024.06.09.597944

**Authors:** Florent Sylvestre, Nadia Aubin-Horth, Louis Bernatchez

## Abstract

Sexual dimorphism can evolve through sex-specific regulation of the same gene set. However, sex chromosomes can also facilitate this by directly linking gene expression to sex. Moreover, differences in gene content between heteromorphic sex chromosomes contributes to sexual dimorphism. Understanding patterns of sex-biased gene expression across organisms is important for gaining insight about the evolution of sexual dimorphism and sex chromosomes. Moreover, studying gene expression in species with recently established sex chromosomes can help understand the evolutionary dynamics of gene loss and dosage compensation. The three-spined stickleback is known for its strong sexual dimorphism, especially during the reproductive period. Sex is determined by a young XY sex chromosome pair with a non-recombining region divided in three strata, which have started to degenerate. Using the high multiplexing capability of 3′ QuantSeq to sequence the sex-biased transcriptome of liver, gills and brain, we provide the first characterization of sex-specific transcriptomes from ∼80 stickleback (40 males and 40 females) collected from a natural population during the reproductive period. We find that the liver is extremely differentiated between sexes (36% of autosomal genes) and reflects ongoing reproduction, while the brain shows very low levels of differentiation (0.78%) with no functional enrichment. Finally, the gills exhibit high levels of differentiation (5%), suggesting that sex should be considered in physiological and ecotoxicological studies of gill responses in fishes. We also find that sex-biased gene expression in hemizygous genes is mainly driven by a lack of dosage compensation. However, sex-biased expression of genes that have conserved copies on both sex chromosomes is likely driven by the degeneration of Y allele expression and a down-regulation of male-beneficial mutations on the X chromosome.

## Introduction

Species with sexual reproduction often exhibit sexual dimorphism (Lande, 1980). Because sexes share most of their genetic material except for potential sex chromosomes, sexual dimorphism can arise from the different regulation of the same set of genes (Ellegren & Parsch, 2007; Tosto et al., 2023). Sex-biased gene expression has been described in many systems as dependent on both life stages (Djordjevic et al., 2022) and tissues (Rodríguez-Montes et al., 2023). Compiled data for five tissues in five mammals and a bird species show that sex-biased gene expression varies in intensity across tissues, species and sexual maturity, with the contribution of sex chromosomes being also variable (Rodríguez-Montes et al., 2023).

The variety of sex determinism systems and reproductive behaviors present in teleost fish make them an interesting vertebrate group to study sex-biased gene expression (Thresher, 1984; Devlin & Nagahama, 2002; Kobayashi et al., 2013). Previous studies have highlighted a wide variation in patterns of sex-biased expression among species and between tissues within species. For example, patterns of sex-biased gene expression have been described in the brain or liver, which are known to play a role in reproduction. Sex-biased gene differentiation in the brain seems highly species dependent, with about a thousand genes identified in salmonids (Hale et al., 2018) but only a handful in the Gulp pipefish *Syngnathus scovelli* (Beal et al., 2018) or the zebrafish *Danio renio* (Yuan et al., 2019). In cichlids, gene expression levels in the brain are associated with social status and gonadic sex (Renn et al., 2008; Schumer et al., 2011). Liver is sexually dimorphic, specifically in oviparous species, where it produces many proteins or proteins precusors which will be stored in the eggs (Qiao et al., 2016; Darolti & Mank, 2023), and many genes have been identified as sexually biased in salmonids (Sutherland et al., 2019) and across cichlid taxa (Lichilín et al., 2021). On the contrary, other tissues such as the gills have received strong attention in the context of adaptation to salinity, pollution or hypoxia (Scott et al., 2004; Van Der Meer et al., 2005; Gonzalez et al., 2006) but studies rarely account for potential sex dimorphism in the response of this tissue (but see Lichilín et al., 2021). Of course, caution should be taken when comparing level of sex-bias across studies with different life stages, protocol, power and methods, which is why we need to characterize variation in sex-biased gene expression across tissue in a single framework in order to better understand it.

Sex chromosomes play an important role in sex dimorphism (Rice, 1984). However, understanding patterns of sex-biased gene expression on sex chromosomes is particularly complex in species with non-recombining sex chromosomes, as they tend to degenerate and may involve dosage compensation (Bachtrog, 2013). The lowered effective population size of non-recombining Y (or W) chromosome, present in only one sex, leads to lowered efficiency of natural selection and the degeneration of Y chromosomes (Charlesworth & Charlesworth, 2000). Genes that have lost their Y or W copy (hemizygous genes) are expected to exhibit a reduced expression level that does not come from sex-specific gene regulation. However, lowered gene activity on sex-chromosome can have widespread effects on autosomal genes (Wijchers et al., 2010), and mutation reestablishing the ancestral level of expression in the heterogametic sex is expected to be advantageous, leading to the evolution of dosage compensation. Global dosage compensation, where the X (or Z) chromosome is overexpressed to compensate for the loss of Y (or W) genes was long thought to be necessary in systems with degenerated sex chromosomes, but accumulating literature outside Drosophila and mammals suggest that it is not necessarily the case (Mank et al., 2011; Mank, 2013), with many groups exhibiting no or partial dosage compensation. Moreover, many genes are dose insensitive (i.e., their copy number does not affect their expression level or protein abundance), therefore they do not need to be compensated. In fish, global dosage compensation has rarely been found when studied (Darolti et al., 2019) and we still lack knowledge about the extent of the evolution of dosage compensation in this highly diverse group.

The three-spined stickleback is a model fish species in behavior and evolutionary biology (Reid et al., 2021) with a young XY sex-chromosome system, at most 13–26 million years-old (Peichel et al., 2020), and a strong sexual dimorphism over colouration, size and behavior during the reproductive period. Reproduction is costly for males, as they build a nest for the female to lay eggs, which they protect from other individuals that might raid it. They recruit females using a specific nuptial parade, after which they fertilize the eggs. Parental care is also handled by males, involving several specific behavior such as nest and eggs cleaning, defense, and fanning (FitzGerald, 1993), activities that are energetically costly (FitzGerald et al., 1989; Chellappa & Huntingford, 1989; Dufresne et al., 1990). Additionally, food intake is reduced in males during this period (Wootton, 1976). Males are smaller but exhibit a red coloration and blue eyes (Barker & Milinski, 1993; Folstad et al., 1994) corresponding to an honest signal for female preference, which coupled with nest-care behavior increases predation risk in comparison to less red males and females, which are green-grey (Whoriskey & Fitzgerald, 1985; Johnson & Candolin, 2017). They also exhibit differences in the prevalence of parasites (Reimchen, 1997; Reimchen & Nosil, 2001) and in general morphology (Kitano et al., 2007).

Sexual dimorphism also exists at other phenotypic levels, and studies have found sex-specific splicing and protein expression (Viitaniemi & Leder, 2011; Naftaly et al., 2021). Sex-biased gene expression has been described from laboratory raised, non-reproductive adults, in the liver (Leder et al., 2010), brain (Kitano et al., 2020; Kaitetzidou et al., 2022), gonad, and adipose tissue (Kaitetzidou et al., 2022). In the liver, Leder et al. (2010) identified that 11% of expressed genes were sex-biased, 24% of which were located on sex-chromosomes, and the rest rather homogeneously distributed across chromosomes. Female-biased genes show enrichment in ribosomal activities, translation and intracellular processes, while male-biased gene are more involved in signaling. In the brain, the two available studies in adults found drastically different results, with Kitano et al. (2020) identifying thousands of sex-biased genes in the adult brain, of which only 8% are located on sex chromosomes. On the opposite, Kaitetzidou et al. (2022) identified less than two hundred sex-biased genes, and 54% of them were located on sex chromosomes. Furthermore, a study in 1-year old lab-reared immature sticklebacks found large differences in gene expression in the brain between sexes prior to gonadal development (1255 genes), with 38% of them being genes found on sex chromosomes (Metzger & Schulte, 2016). Gonads unsurprisingly exhibited strong sex-biased expression, with ∼2400 sex-biased transcripts, and adipose tissue was the least differentiated, with only 67 sex-biased genes. However, these studies have relied on laboratory raised, non reproductive populations, and we lack information about how reproduction might affect these patterns.

The sex chromosome consists of a pseudoautosomal region which is still recombining, and three evolutionary strata which have evolved through successive inversion (Peichel et al., 2020). Accumulation of sex-biased genes on sex chromosomes seems to be associated with a lack of global dosage compensation (Leder et al., 2010; Schultheiß et al., 2015; White et al., 2015) in that species, coupled with a potential partial dosage compensation in stratum I of the Y chromosome. While these studies provide an interesting portrait of sex-biased gene expression across tissues, they often use different statistical approaches, which makes them difficult to synthetize. We need more studies similar to Kaitetzidou et al. (2022) comparing tissues in a single methodological and analytical framework to better understand the extent of inter-tissue variation in sex-biased gene expression. Furthermore, and most importantly, we still lack knowledge about sex-biased gene expression during the reproductive period, which corresponds to the expression of the sexually dimorphic characteristics of the three-spined stickleback.

In this study, we took advantage of the high sample multiplexing capability offered by QuantSeq 3′-UTR sequencing (Moll et al., 2014) to profile the transcriptome of ∼40 samples per sex in three tissues (liver, brain, and gills) in adults from a natural population of three-spined stickleback from eastern Canada. We aimed at 1) describing the sex-specific transcriptome of this species in each tissue and 2) making use of the recently sequenced Y chromosome data to understand the dynamics of dosage compensation. According to work in other species and stickleback, we expected the liver to be a highly differentiated somatic tissue between sexes, as it plays an important role during reproduction in teleost fish (Leder et al., 2010; Meng et al., 2016). We also expected the brain to be strongly sex-biased, as the differences in reproductive behavior between sexes are striking, and the tissue itself is known to be sexually dimorphic during the reproduction of the three-spined stickleback in term of size (Kotrschal et al., 2012). Transcriptomic work also suggested strong differentiation (Kitano et al. 2020) but not always (Kaitetzidou et al., 2022), making predictions difficult. Finally, sex is not a factor usually accounted for when studying gills, an extremely important tissue for physiological regulation, and when it is, there is almost no sex-biased expression (Lichilín et al., 2021). We therefore predicted that this would be a neutral tissue with little to no sex-biased expression. We also expected to find that most genes on sex-chromosome exhibit sex-biased gene expression caused by the lack of global dosage compensation in that species.

## Materials and Methods

### Ethics statement

This study was approved by the Comité de Protection des Animaux de l’Université Laval (CPAUL, approval number SIRUL 109096). A fish permit was issue by the Ministère des Forêts, de la Faune et des Parcs du Québec (permit number 2018 04 11 005 01 S P) for sampling.

### Sampling and Sequencing

We collected adult anadromous three-spined sticklebacks from tide pools of the St. Lawrence River at Baie de l’Isle verte (48.009961, −69.407070) in July 2018. Brain, liver, and gills were dissected (under five, seven and ten minutes after death respectively) and preserved in RNAlater at −20 C. In 2022, we performed RNA extractions using the RNeasy mini-Kit (Qiagen). We disrupted samples in 700 μL (brain) or 1400 μL (Gills, Liver) of trizol using a mixermix 400 at 30 Hz for 3 minutes or until complete tissue disruption. After three minutes, we added 140 μL (or 280 μL) of chloroform, homogenized the solution and waited five minutes before centrifuging for 15 min at 12,000g. We collected the upper phase and added 550 μL (or 1100 μL) of ethanol before transferring 700 μL into a RNeasy Mini column (Qiagen). We then centrifuged for 15s at 11,000g, discarded the flow-through and repeated the operation until all the collected phase has been used. We then proceeded with the extraction protocol following manufacturer’s instruction, including a DNase step, but replacing buffer RW1 by buffer RWT from miRNeasy Mini Kit (Qiagen), as it yielded better results for the brain. We then generated QuantSeq libraries using 3′ mRNA-Seq Library Prep Kit (Lexogen) with dual indexing to identify each individual and 18 PCR cycles for library amplification. QuantSeq 3’UTR library preparation allows to quantify gene expression by amplifying only the 3’ end of each transcript using poly-A tail priming. This approach reduces the required coverage for estimating gene expression levels but removes the possibility of study alternative transcript and sequences variation, allowing the study of many samples at a reasonable cost. After quality check on a 2100 Bioanalyzer (Agilent Technologies) and concentration estimation using Quant-iT PicoGreen (Invitrogen), 227 libraries were pooled to equimolarity and sent to the Centre d’Expertise et de Services Génome Québec (Montréal, QC, Canada) for 50 bp single-end sequencing on 2 lanes of an Illumina Novaseq X. A read length of 50 bp was selected as longer sequencing will only result in longer sequencing of the poly-A tail, which does not contain helpful information about the gene sequence. Tissue disruption and quality check were performed at the Plateforme d’Analyse Genomique of Université Laval (Québec, QC, Canada).

### Alignment and Expression Counts

After quality check using FASTQC v0.11.8, we used fastp v 0.15.0 (Chen et al., 2018) to trim poly-A tails, Illumina adapters, and the first 12bp of each read, as they showed biased composition. We discarded reads shorter than 20bp long and proceeded with the alignment. We used STAR v2.7.2b (Dobin et al., 2013) two-pass mode to align reads to the stickleback reference genome V5 (Peichel et al., 2020; Nath et al., 2021), accessible on NCBI as RefSeq assembly GCF_016920845.1), excluding the Y reference sequence for the females and its pseudo-autosomal region (PAR) for males. We used STAR two-pass mode to discover reads junctions and improve mapping accuracy in the second pass. We then quantified gene expression using HTSeq v0.11.3. (Anders et al., 2015) using htseq-count in union mode and no strand constrain (−s no) after filtering out multi-mapping reads using samtools (Li et al., 2009). Counts from each sample were merged using custom R scripts, leading to four datasets: the total datasets comprising all samples, and one dataset for each tissue (Liver, Gills and Brain).

### Filtering, normalization, and quality check

Unless stated otherwise, analyses were performed in R v 4.3.2 (R Core Team, 2021) and python 3.9.12 (Rossum & Drake, 2010). For each dataset, we applied the same filtering procedure. First, samples with fewer than 5,000,000 raw reads counts were excluded, as preliminary principal component analysis (PCA) showed that they clustered together (result not shown). Then, we kept only genes with one count per million (cpm) in at least 10 samples. Cpm normalization allows comparison of gene expression level by normalizing read counts by the total number of reads per sample. To identify potential errors in the dataset (in particular, whether tissue was misidentified for some samples), we first performed a PCA on autosomal genes across all samples, using blind variance stabilization transformation (Vst) as implemented in DESeq2 v1.40.2 (Love et al., 2014), which normalize for the increases in variance with mean gene expression, and is recommended for clustering-based analysis. For all other analyses, read counts normalization was carried out independently for each tissue using the median of ratio normalization factor implemented in DESeq2, which account for differences in number of reads and gene composition of each sample, to allow precise comparison of gene expression across samples. Given the specificity of genes on sex chromosomes, only autosomal genes were used for the calculation of the scaling factors in each tissue, which were then used to normalize sex-chromosome genes independently. Note that gene length was not accounted for in the normalization process, as it could generate a bias in our dataset as QuantSeq only sequences the poly-A tail of each transcript, not the full transcript. Additionally, we identified genes representing more than 10% of total read counts in any individual (hereafter named “highly expressed genes”).

### Differential Expression Analysis of Autosomal Genes

Within each tissue, we used Wilcoxon rank-sum tests to compare gene expression between sexes and identify differentially expressed genes (DEG), as suggested by a recent study, which showed that the classically used negative binomial models implemented in Deseq2 are subject to increased false positive rates in large sample size datasets (Li, Ge et al., 2022). We used the Benjamini-Hochberg procedure to control the false discovery rate (Benjamini & Hochberg, 1995), using a 5% q-value for significance. We also calculated the log-fold change (LFC) of gene expression between sexes as, for each gene:

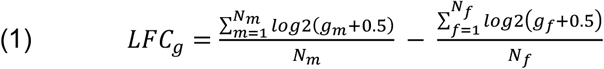

with gm and gf being the normalized read count for each male and female, and Nm and Nf the number of individuals from each sex.

### Functional Characterization of Differentially Expressed Genes

To explore common functions among sex-biased genes, we performed Gene Ontology (GO) enrichment analysis. To do so, we used blastX (Camacho et al., 2009) to gather the Swiss-Prot (Schneider et al., 2009) annotation for each sequence of the three-spined stickleback transcriptome available on NCBI (RefSeq assembly GCF_016920845.1) and gathered gene ontology information from the associated UniProt entry (The UniProt Consortium, 2023). We then summarized transcript-level GO at the gene level using information from the NCBI annotation of our reference genome with a custom python script. We used goatools (Klopfenstein et al., 2018) to perform Fisher’s exact test for enrichment at a q-value threshold of 0.05 using the Benjamini-Hochberg procedure, using the set of expressed genes in each tissue as the reference set, to distinguish the enrichment of sex-biased genes with enrichment driven by the specific role of each tissue. We also combined Zfin (zfin.atlassian.net), genecards (www.genecards.org) and Uniprot (www.uniprot.org) databases to provide a more precise functional characterization of 1) the 10 genes with the lowest q-value and 2) the 10 genes with the highest LFC for each tissue, as well as all DEG shared by all tissues. Functional annotations described in the results section are based on information from these databases unless otherwise noted.

To explore the genomic distribution of DEGs, we performed a Fisher test for enrichment analysis for each chromosome to test whether 1) some chromosomes are enriched in DEG considering their number of genes and 2) DEG within a chromosome are enriched toward male-biased or female-biased genes considering the global distribution of sex bias toward each sex. We used a Benjamini-Hochberg procedure to control the false discovery rate independently for the two hypotheses tested.

### Identification of Shared Genes Between X and Y

The study of sex-biased gene expression on sex chromosomes is complicated by independent gene gain or loss on their non-recombining region. To identify gene loss or gain, we first extracted transcript sequences from the reference genome using gffread (Pertea & Pertea, 2020) and the NCBI genome annotation for our genome version, using -C option to remove transcripts with no CDS. We did the same using the available nine-spined stickleback (*Pungitus pungitus*) reference genome (Varadharajan et al., 2019) and annotation (GenBank accession GCA_949316345.1), which we use to better infer the relationships (orthology or paralogy) between genes. We then used Orthofinder v2.5.5 (Emms & Kelly, 2019) to identify orthogroups with default parameters. Orthofinder uses reciprocal best hit between transcript sequences to infer orthology relationships between them. The result is a file presenting, for each species, the list of transcripts belonging to the same group, which can be used to infer orthology between species as a group of sequence with only one member in each species. However, because we are interested in the relationship between two chromosomes within the same species, our interpretation differs in that orthology between X and Y corresponds to a group or at least two sequences from the same species. Additionally, the presence of several transcripts per gene further complicates the matter, as all transcripts from the same gene are expected to cluster together, leading to groups of more than two sequences still corresponding to orthologs. We used a custom python script to identify transcripts likely to be orthologs between X and Y chromosomes and summarize the information at the gene level, using information from the NCBI genome annotation for the three-spined stickleback, which contains independent annotation of X and Y chromosomes. Orthologs between X and Y chromosomes were defined as an orthogroup composed of one transcript from X, one from the Y and other members originated from the nine-spined stickleback. To account for the fact that one gene can have several transcripts, we accepted an orthogroup with many X or Y transcripts if they belonged to the same X or Y gene. However, we observed that for some genes, not all transcripts clustered in the same orthogroup, and removed such genes from our analysis, except in the situation were most transcript clustered together, and one transcript wasn’t assigned any orthogroup. Transcripts that fell in an orthogroup with only Y or X transcripts were categorized as hemiploid Y (no gene copy on X) or hemiploid X (no gene copy on Y). Orthogroups that contained autosomal genes, likely reflecting gene gain on the sex chromosome were removed from analysis as well as genes with multiple copies on either sex chromosome. Finally, genes for which transcripts couldn’t be assigned to any orthogroup were considered as hemiploid X or Y.

Gene counts for genes still shared between X and Y chromosomes were pooled in males. We assigned each gene to one of the three known evolutionary strata of sex chromosomes using its central position, and breakpoints defined in Peichel et al.(2020). Given that the pseudoautosomal region of the Y chromosome was excluded from the read mapping step, read counts for that region were already correct and genes were considered as still sharing a copy, ignoring Orthofinder results.

### Sex-Biased Gene Expression on the Sex Chromosomes

We used the merged counts to test for sex-biased gene expression on sex chromosomes using the same method as for the autosomes. To test for potential dosage compensation, we estimated the autosome to sex chromosome median expression ratio for all genes, and independently for hemiploid X and Y genes for males and females. The autosomal median was calculated as the median across all genes of the mean log2 normalized read counts across all samples. Median expression for genes on the sex chromosome was calculated similarly but separating individuals by sex. We estimated one median for all genes, genes that still have a copy on both chromosomes and genes specific to each sex chromosome. This calculation was done for the whole chromosome and for each stratum independently (PAR, strata I, strata II and strata III). Confidence intervals (CI) were calculated using bias correction bootstrapping for both autosomal and sex chromosome genes and significance assessed using overlap of the CI with 0. Finally, to understand the processes underlying the evolution of sex-biased gene expression for genes with a copy conserved on each chromosome, we estimated X and Y allele expression in males. We then compared the expression level of female, male and X and Y alleles in sex-biased genes to their expression levels in unbiased genes using Wilcoxon rank-sum test.

## Results and discussion

### Sequencing, Mapping and Filtering

The two lines combined rendered 4,247,758,804 reads with an average of 20,319,408 [CI 95%: 18,047,379 – 22,591,436] for the gills, 18,847,630 [17,937,603 – 19,757,656] for the brain and 17,073,329 [14,606,317 – 19,540,341] for the liver. The average read length after quality trimming was 38 bp. After removing samples with fewer than 5,000,000 reads (ten for the liver, one for the brain), we kept 67 samples in the liver (35 Females; 32 Males), 77 for the brain (38; 39) and 72 in gills (36; 36). The dataset included 59 samples sequenced in all tissue, but 10 were specific to gills and brain, five to brain and liver and three to gills and liver. Three samples were sequenced only in the brain. We had a percent mapping of uniquely mapped reads to the reference genome ranging from 44.3% to 88.6% (median 76.8%). Unmapped reads in low quality samples mainly corresponded to ribosomal RNA and external transcribed spacer, genes known to have multiple copies across the genome. Correlation between mean expression levels estimated on the whole dataset and in a subset of 10 males and 10 females between 70-80% of mapping rates were all above .99%, suggesting that our normalization process was effective in correcting the differences in total coverage in samples with low mapping rates. After filtering, we identified five overexpressed genes in the liver but none in the gills or the brain. Liver was the tissue with the lowest number of expressed autosomal genes (14,402) compared to gills (17,624) and brain (17,930), with 13,957 autosomal genes expressed in all three tissues. Using a PCA to screen our dataset for potential cross-tissue contamination or mislabelling revealed no evident issue (Fig.1A), as each tissue formed a distinct group, confirming the quality of the dataset.

**Figure 1:**
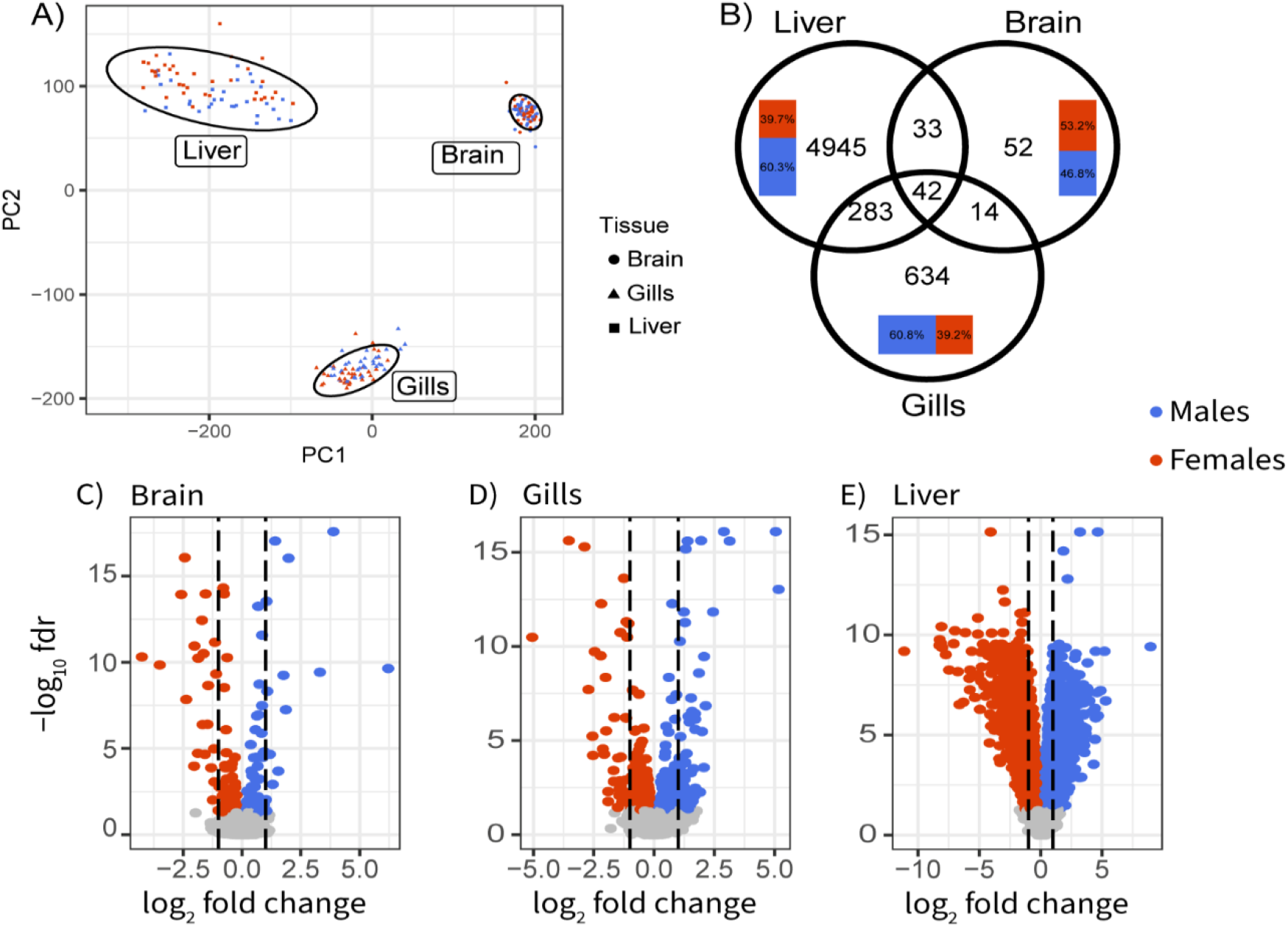
Patterns of autosomal sex-biased gene expression in three-spined stickleback. A) PCA of gene expression on all tissues. B) Overlap of sex-biased genes between tissues. Inset bar plot represents tissue specific repartition of male (blue) and female-biased genes. Volcano plots of sex-biased gene expression in C) brain, D) gills and E) liver. Colored dots correspond to genes significant at a 5% false discovery rate based on a Wilcoxon-rank-sum test. Dotted lines represent a log-fold change of 1 (doubled expression).

### Tissues Differ in the Magnitude of the Sex-Biased Expression

Patterns of sex-biased gene expression varied greatly between liver, gills and brain, both in terms of the number of genes differentially expressed and their function (Fig, 1B). Liver transcriptomes showed strong sex specificity, with 5,303 sex-biased genes (hereafter SBG, “sex-biased gene”) with a q-value ≤ 5% after a Benjamini-Hocheberg correction. In comparison, using the same threshold, we found 973 SBG in gills and only 141 in the brain (Fig. 1B). Genes also exhibited a wider range of differential expression in the liver compared to gills and brain (Fig. 1C, D, E). Across all tissues, no chromosome showed enrichment for SBG (Fig. 2, Fisher’s test q-value> 0.4). All SBG are listed in Table S1.

**Figure 2:**
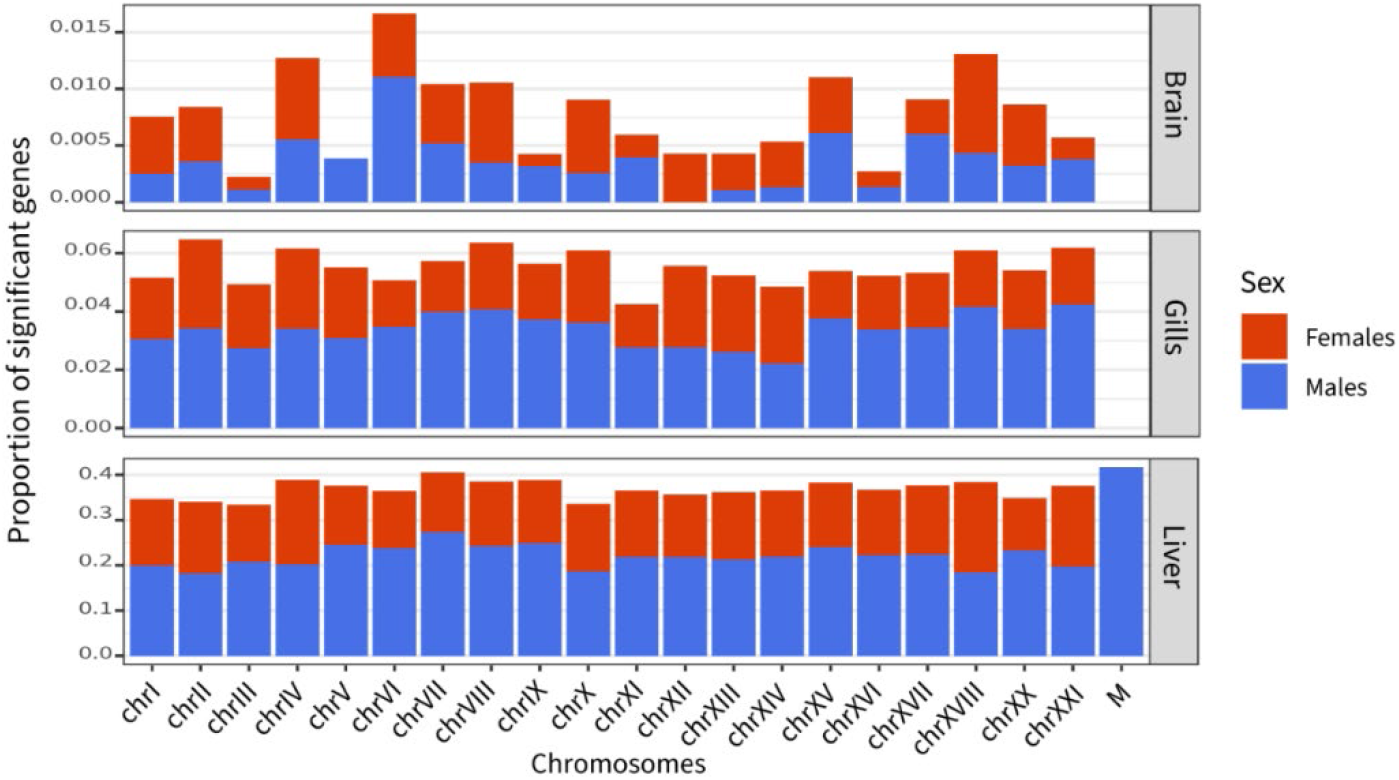
Proportion of sex biased genes across chromosomes and sex in each tissue. Chromosome IV and VII show enrichment toward male-biased function in liver (q-value ≤0.05), and chromosome XVIII toward females-biased functions as well as the mitochondria with lower confidence (q-value = 0.12. No chromosomes are significantly enriched in sex-biased genes.

### Sex-biased genes found in all tissues are implicated in cell physiology, cell-cell signalling and gene expression modulation

Most SBG were unique to a tissue, with only 42 significant genes overlapping in all three tissues (Fig. 1B, Table S2). Eight of them were related to basic cell physiology such as growth, cytoskeleton, and differentiation. Five genes were involved in cell-cell signaling or adhesion and are mainly known to play a role in neuron communication and development. Seven genes were involved in gene expression modulation and affect either DNA methylation, transcription, or alternative splicing. In fish, sex-specific methylation is known for its role in modulating the expression level of reproduction-related genes (Laing et al., 2018; Li, Chen, et al., 2022) and in sex determinism (Gemmell et al., 2019). Seven genes were involved in gene expression modulation and affect either DNA methylation, transcription, or alternative splicing. In fish, sex-specific methylation is known for its role in modulating the expression level of reproduction-related genes (Laing et al., 2018; Li, Chen, et al., 2022) and in sex determinism (Gemmell et al., 2019).

Similarly, sex-specific alternative splicing can provide an alternative route from gene regulation to generate sexual dimorphism (Telonis-Scott et al., 2009; Naftaly et al., 2021). Hence, those genes could play an important role in regulating sex-biased gene expression and dimorphism across tissues. Other functions found in shared SBG among tissues involve the immune system, testosterone response (one gene) and two nuclear genes with mitochondrial function.

In most cases, genes showed the same directionality in all tissues (Fig. 3), except for two genes: slc16a13, a monocarboxylic acid transporter, and esr2b, an estrogen receptor. Both are female-biased in liver but male-biased in the brain and gills. While the function of slc16a13 is hard to interpret in our context, as the substrate of this member of a large family of solute transporter is unknown (Halestrap, 2012), the gene esr2b code for an estrogen receptor. In teleosts, estrogen receptors are involved in several biological processes, including reproductive processes (Nelson & Habibi, 2013). In the brain, esr2b is thought to play a role in reproduction through the regulation of gonadotropin production, which is involved in gametogenesis (Muriach et al., 2008) in both sexes and have been found to have a higher expression in males in the pituitary gland of the fathead minnow (Pimephales promelas) (Filby & Tyler, 2005). In the liver, esr2b showed strong expression levels and was female biased (LFC of - 0.35), and could be associated with vitellogenesis process, although it seems to vary across species (Dominguez et al., 2014; Chen et al., 2019), which prevent us to interpret it in the context of the three-spined stickleback. We still lack knowledge on the effect of sex hormones in the gills.

**Figure 3:**
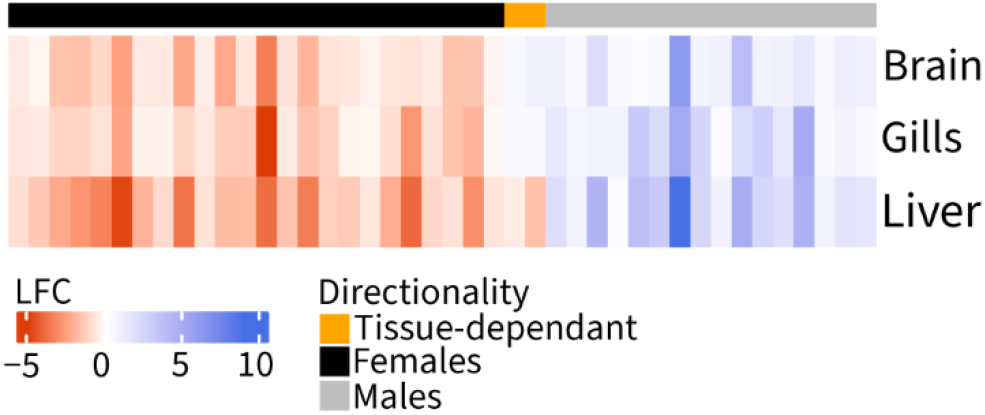
Heatmap of log-fold change in gene expression between sexes for 42 genes significantly differentially expressed in all tissues. Bar on top shows concordance in the direction of expression bias.

In the case of a pattern of directionality in gene expression shared by only two tissues, we only found discordance when the comparison included the liver (Fig. S1). While part of this is explained by the liver having both more SBG and more shared SBG with other tissues in general, it suggests that this tissue might have a different usage of the same gene set. These results are in accordance with other studies, which find that sex-biased genes are often tissue-specific (Mank et al., 2008; Mayne et al., 2016; Rodríguez-Montes et al., 2023). The comparative analysis of several tissue highlights that while a core set of genes probably plays a central role in sexual dimorphism throughout the body, potentially by regulating other key processes, the study of multiple tissues is important to understand the extent of sexual dimorphism, and to understand the forces driving the evolution of sex-biased gene expression.

### Sex-biased genes in the brain are few and not enriched for particular functions

The brain showed the lowest number of SBG, with 141 genes differentially expressed between sexes on the autosomes (0.78% of expressed genes), equally distributed between males and females and across chromosomes (Fig. 2). We found no enrichment for any gene ontology term using the 5% threshold. Looking into the most significant genes (10 with highest p-value and 10 with highest LFC), we found that only 12 of them are specific to the brain (Table S3). We were unable to annotate three of them (LOC120812970, LOC120817963 and LOC120833148). The remaining 9 genes were associated with various biological functions. The growth hormone-releasing hormone (ghrh), more expressed in males, is the first hormone secreted in the growth hormone axis, which in fish is not only involved in growth but also reproduction, metabolism and immune function (Chang & Wong, 2009). TTC29, also more expressed in males, is involved in cilium movement and is mainly described in sperm flagellum (Bereketoğlu et al., 2022). Other genes more expressed in males included ihhb (LOC120819658), which is involved in neural and chondrocyte development (Wu et al., 2001; Chung et al., 2013), and MMP13 or 18 (LOC120822795), which modulates angiogenesis in the brain (Ma et al., 2016). In females, significantly biased expressed genes included hcn3 (si:dkey-197j19.6), an ion channel which is essential for neuronal function, nlrc3 (LOC120810501), which has been implicated in neuromast cell regeneration in zebra fish (Jiang et al., 2014), and ecm1a (LOC120810788), which codes for an extracellular matrix protein involved in signal transduction.

Overall, the differences between the sexes were small for this tissue and implicated various functions associated with neural tissues, without clear divergence between males and females. These results are in the same order of magnitude than the ones from Kaitetzidou et al. (2022), which identified 104 sex-biased autosomal genes in the brain. Results strikingly differ from those of Kitano et al. (2020), which categorized 59.3% of all genes to be sex-biased in the brain (∼7000 genes). Methodology differs, as they did not used multiple tests correction to detect as many sex-biased genes as possible, but if we use a similar threshold (p-value <= 0.05) on our dataset, we still only identify 1289 significant genes. Differences might also stem from the number of samples, our dataset allowing for more precise estimates of sex-bias. On the other hand, working in an environmental setting with multiple classes of individuals, such as different age classes and reproductive states might increase variance in gene expression in both males and females compared to laboratory condition. Our field approach has the potential to generate differences but also to hide small differentiation in noise, which complicates the comparison. Furthermore, the extent of sexual dimorphism differs between populations of three-spined sticklebacks (Kitano et al., 2012), at least for morphology, which could be a factor here, although we do not have direct measurements to characterize the extent of differentiation in our population. Finally, our results are in contrast with the level of dimorphism shown in behavioral and morphological studies (FitzGerald, 1993; Kotrschal et al., 2012; King et al., 2013). An explanation lies in the timing of development of these differences in the three-spined stickleback and other fishes. It is known that differences between the sexes in this species are governed by hormonal and neuropeptide changes, whose levels already diverge prior to the actual reproduction period and are directly linked to adult sexual dimorphism (Petersen et al., 2015). This can be exemplified by affecting hormonal levels during a critical developmental window in embryonic sticklebacks, by exposure to an endocrine disruptor, which results in significant changes in androgen levels already at the larval stage, which in turn affects the hormonal reproductive axis and gonadal development in adults (Petersen et al., 2015). The onset of sexual maturation is controlled by photoperiod in this species and the different neuropeptides and hormones that are activated do so at different time intervals. For example, some are significantly differentially expressed after 10 days once males are exposed to a long day /short night photoperiod, while others take 30 days to increase significantly (Shao et al., 2019). Furthermore, once mature, behavior and hormonal levels differ between different phases of reproduction, such as courting and nesting in males (Mayer et al., 2004; Kent & Bell, 2018) and the different stages of egg production in females(Graham et al., 2018). It is possible that when we sampled wild individuals, we captured these within sexes differences, in both sexes, affecting the number of observed differentially expressed genes in the brain, whose quantified fold changes are lower than other tissues studied here and thus for which we have lower statistical power.

### Sex-biased genes in gills are associated with ion-related functions and immune defense

We identified 973 SBG in the gills (5.5% of expressed genes), of which 60.8% were male-biased. Significant genes were distributed homogeneously in the genome, with no chromosome significantly enriched in SBG, nor SBG enriched in males of females biased genes or biased toward a sex (Fig. 2). Synaptic signaling and organization represented 48% of significantly enriched GO terms (Table S4). However, looking at descriptions of genes within those GO categories on Zfin or Genbank revealed that many genes code for ion channels or pumps, which are a core function of the gill tissue (Perry et al., 2003), yet the associated GO functions have only been described in the brain (Table S5). This suggests that gene ontology analysis in gills suffer from the lack of gill-specific information. Other GOs included various biosynthetic and metabolic processes, as well as cell adhesion, communication, and development. The most strongly differentiated genes (Table S6) in the gills involved two genes with potential role in pathogen resistance, hhipl1 and CLEC4M (LOC120817010), both more expressed in females. Asic2, also biased toward female gills, codes for an ion channel, and this gene family is involved in Na+ intake in rainbow trout gills (Dymowska et al., 2014), while si:dkeyp-92c9.4 has no specific function uncovered. Finally, the galanin receptor galr2b, also biased toward females, is associated with adenylate cyclase-modulating G protein-coupled receptor signaling pathway and neuropeptide signaling pathway. In males, the most differentiated genes (Table S6) were associated with basic cellular functions (LOC120828377, and LOC120810538). Two genes in males were involved in neuronal function, including one ion channel, rem2, and Dpysl3, also male-biased in the liver, which is proposed to have an effect of peripheric axon growth. Finally, three genes (LOC120816929, LOC120817829 and LOC120821053) could not be annotated.

Most studies on gills transcriptomes focused on their role in osmoregulation and respiratory processes in responses to anoxia, salinity, or various contaminants (Scott et al., 2004; Van Der Meer et al., 2005; Gonzalez et al., 2006). Works have also been interested in gills’ function in defense against pathogens, as they represent a direct entry for infection and parasites (Mitchell & Rodger, 2011). However, a survey of the physiology literature illustrates that these studies do not usually account for sex. When they do, few SBG are identified, for example, in wild populations of African cichlids (Lichilín et al., 2021). However, our results identify numerous SBG between sexes, all found in the same environmental conditions, with 60% of genes more expressed in males. Sampling during the reproductive period, which is costly for both sexes and in particular in males, might explain the differences observed with previous work. This unexpected result highlights the importance of accounting for sex when studying the gills, as the extent to which this tissue might respond differently to various challenges between sexes is also poorly understood.

### The Liver is a Hotspot of Sex-Biased Gene Expression

We identified 5,303 SBG in the liver (36.8% of total expressed genes), 60.3% of which were male-biased. SBG were uniformly distributed among chromosomes (Fisher’s exact test qvalue ≥ 0.45, Fig. 2), except for chromosomes IV and VII, which showed enrichment in male-biased genes (FDR ≤ 0.05), XVIII, toward female-biased genes (FDR ≤ 0.05) and the mitochondria which showed marginal enrichment for male-biased function (FDR of 0.12). This, with similar results previously described in the brain and gills, suggests that the genetic architecture of sexual dimorphism is uniformly distributed across the genome.

While comparing the number of genes between studies is complex as life stages, condition and filtering have a deep impact on detected SBG, widespread sex-biased gene expression has been found in the liver of other fish species, such as lab-raised *Salvelinus fontinalis*, in which SBG represent 16.1% of the total gene expression using more stringent filtering (LFC>= 1.5) than we applied (Sutherland et al., 2019). Similarly, very low levels of SBG are identified in cichlids (Lichilín et al., 2021) using a LFC of 2 as a cutoff but these observations vary across species. In our study, only 396 (2.6%) in the liver passed this cutoff (644 for the 1.5 cutoff), suggesting that sex-specific regulation of gene expression in the liver mostly occurs through subtle regulatory changes, with a median absolute LFC for significant genes at 0.63. Enriched gene ontology terms in the liver were mainly related to metabolic and biosynthetic processes. The immune system also seemed differentiated between the sexes, with enriched processes such as humoral immune response and response to external stimulus. We also identified an enrichment in hemostasis and coagulation regulation (Table S7). These results are in line with previous studies in the liver (Qiao et al., 2016; Sutherland et al., 2019), which confirm the role of the liver as a strongly sexually dimorphic tissue in teleost fishes.

Among the 25 most significant genes in terms of p-value and fold change (Table S8), numerous genes that were female-biased were involved in response to estrogen and estradiol (fam20cl, vtg3, and LOC100190882, LOC100190880, LOC120823934, blasting to vtg1, and two vtg2 respectively). Other genes showed functions that were found in both sexes, related to gene expression regulation (e.g. lbx2 higher in males, st18 higher in females), response to pathogens (LOC120810467, LOC120820940, blasting respectively to the fucolectin-1, higher in females, and CHIT1, higher in males), as well as cellular differentiation (LOC120824638, blasting to srda3a, which was more expressed in males).

We identified five genes representing more than 10% of reads in at least one sample: apoa2, lect2.1, LOC120808851, LOC120810467 and LOC120823934, hereafter referred to as highly expressed genes (Fig. 4). All five highly expressed genes showed sex-biased gene expression, according to Wilcoxon rank-sum tests (p.value <10-6). Apoa2 (apolipoprotein A-II) and lect2.1 (leukocyte cell derived chemotaxin 2.1), which were more expressed in males, have functions related to lipid transport and immune system. Apolipoproteins are involved in lipid transports in vertebrates but have also been found to have antimicrobial activity in teleost fish (Concha et al., 2003), among other functions. Leukocyte cell derived chemotaxin2 have been known to have chemotactic activity in humans, but also have antibacterial activity in other vertebrates and in teleost fish. Lipid management seem to be a crucial function for males during the reproductive period, as liver energetic reserves (lipids and glycogen) have been show to drop in reproductive males (Chellappa & Huntingford, 1989), as a result of the strong energetic expenditure and reduction in food intake associated with reproductive behavior in males (Wootton, 1976). BlastX results for female-biased genes (LOC120808851, LOC120810467 and LOC120823934, figure 4) are indicative of ongoing egg production in females. LOC120808851 is located on the sex chromosome and shows similarity to the ZP3 (Zona pellucida sperm-binding protein 3) Uniprot entry, a protein that mediates sperm-binding during fertilization. According to Orthofinder results, it is part of a cluster of duplicated genes on the chrXIX (LOC120808849, LOC120809240, LOC120808850, LOC120808851) with a single copy conserved on the Y chromosome, which supports the importance of this function for females. LOC120810467 shows similarity to the Fucolectin-1 entry, which belongs to a family of genes involved in innate immunity (Honda et al., 2000) that has been found to be accumulating in European seabass’ (Dicentrarchus labras) eggs (Parisi et al., 2010). Finally, LOC120823934 shows similarity to Vitellogenin-2, which is a precursor to several egg-yolk proteins (Tata, 1976). This result is concordant with the observation of high levels of estrogen receptors in liver we observed, and further confirmed by the presence of vtg3, another vitellogenin coding gene, and vtg-2 among the list of most significant genes in liver (LFC = −5.69). As mentioned above, other genes related to response to estrogen are among the most significantly differentially expressed genes, and both ZP3 and vitellogenin are genes known to be expressed in liver (Sano et al., 2017), at least in teleosts, further confirming the quality of the dataset.

**Figure 4:**
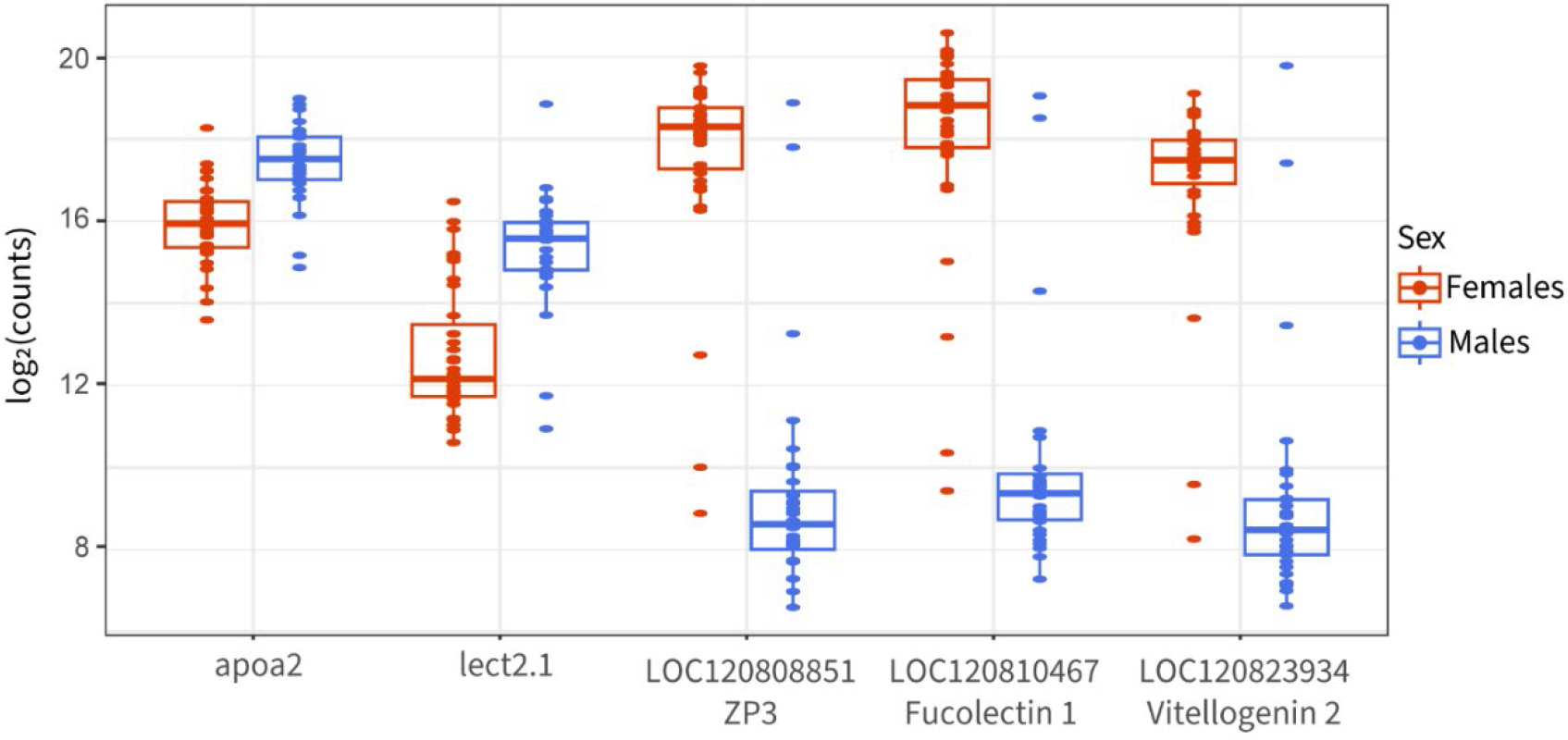
Normalized read counts distribution between sexes for five genes representing more than 15% of total read counts in at least one sample. All between-sex comparisons are significant using an FDR threshold of 5%.

The liver is the only tissue in which we observed sex-biased expression of mitochondrial genes. We identified 13 genes with sex-biased gene expression (∼56% of expressed mitochondrial genes in the liver), all male-biased. These genes included ATP6 and 8, COX2, ND1,2,3,5 and two transfer RNA. In sticklebacks, parental care by the male during the reproductive period, coupled with gonadal development, is associated with a strong depletion of energy reserves (Chellappa et al., 1989; Huntingford et al., 2001). While the development of eggs is also costly for females, the strong involvement in nest building, defence, and parental care by males could generate a high energetic need in males associated with the metabolic processing of energy reserves in the liver. Mitochondrial over-expression thus reflects the strong energetic expenditure of males during the reproductive period.

### Sex-biased Gene Expression on Sex Chromosome Mostly Reflects Gene Loss in Non-Recombining Regions

Most genes located on sex chromosomes exhibited sex-biased gene expression (499 in liver, 755 in gills and 813 in brain, respectively 71%, 83% and 87% of expressed genes on sex chromosomes). Contrasting with the autosomal pattern of sex-biased gene expression, gills and brain exhibited the strongest pattern of SBG, and the higher number of genes expressed in those tissues is not sufficient to explain it. Most SBG are caused by genes having lost their Y copy (84%, 72% and 70% of significant genes in liver, gills and brain), suggesting a lack of global dosage compensation in all studied tissues. Genes orthology relationships for sex chromosomes are available in Table S9. We identified 235 expressed genes still having both their X and Y copy in the brain, 545 having lost their Y copy and 144 their X copy. Numbers are similar in the gills (respectively 242, 528 and126 expressed genes) and liver (204, 405 and 85). Detailled results by strata are presented in Table S10. When looking at the ratio of gene expression between sex chromosomes and autosomes, we found that gene expression was greatly lowered uniquely in males for genes having lost their Y copy (95% confidence interval : [−1.77; −1.29] in brain, [−1.5;−1.00] in gills and [−1.50;−1.01] in liver for males; [−0.71; −0.28], [−0.41; −0.03] and [−0.70; −0.07] in females), but not for genes still having a Y orthologue ([−0.48; 0.24], [−0.59,0.22] and [−0.26, 0.18] in males; [−0.23; 0.39], [−0.27;0.5] and [−0.34, 0.71] in females). This pattern holds across all evolutionary strata and tissue (Fig. 5, Table S10). Estimated ratios for the pseudo-autosomal regions or genes with coding sequences on both chromosomes in non-recombining strata did not statistically differ from 0 (Table S10), except for female-biased expression in stratum II in the brain ([0.10, 1.35], and stratum I in the gills ([0.39, 1.61]). Genes having lost their X copy exhibit a similar pattern but with overall higher median sex-chromosome to autosome expression ratio. In stratum I, it is not statistically different from 0 ([−1.31, 0.02], [−1.44, −0.10] and [−0.92, 0.69]), but is in stratum II and III. As the Y chromosome is likely to conserve or acquire male-beneficial gene, they migth have evolved high expression if they correspond to crucial male function, especially in the I stratum which is the oldest one and have add more time to evolve. Lack of dosage compensation has already been described in the brain (Schultheiß et al., 2015; White et al., 2015) and our results extend this conclusion to the liver and the gills, and to genes with a lost X copy, which had not been included in previous studies. Dosage compensation is expected to evolve when reduced expression in the heterogametic sex affects phenotype, i.e., affects the protein level and its interaction network within the organism. In sticklebacks, conserved genes between the X and Y chromosomes are enriched in haplo-insufficient function (Peichel et al., 2020) and evolving under purifying selection (White et al., 2015), meaning that losing one copy would affect their expression level, potentially affecting their stoichiometry with other cellular component. which might suggest that lost genes were not as impacted by deleterious mutations affecting expression levels. This, coupled with the apparent lack of dosage compensation, suggests that there is no selective pressure to evolve dosage compensation.

**Figure 5:**
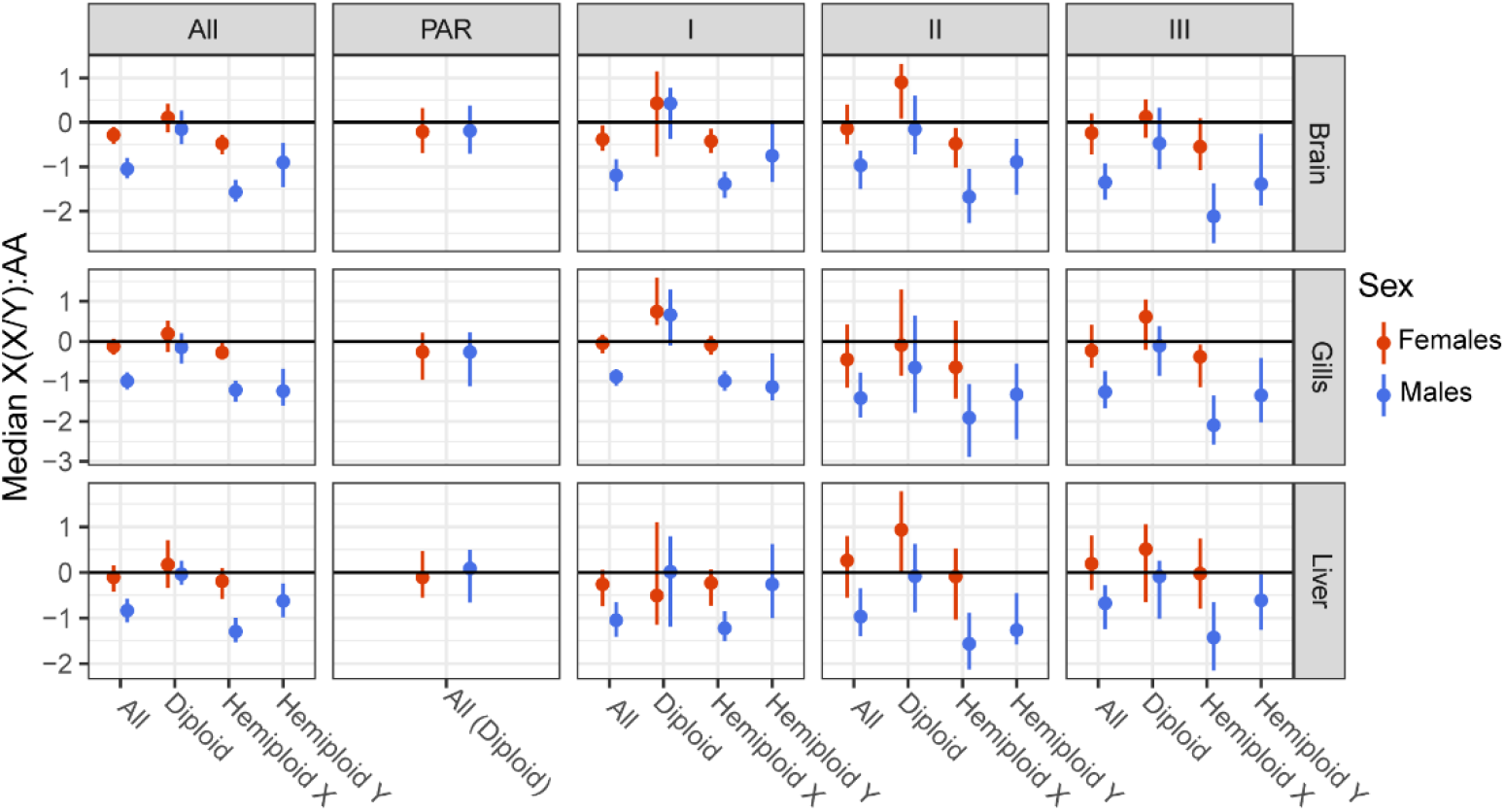
Sex chromosomes to autosome expression ratio across sex-chromosome evolutionary strata. Hemizygous X and hemizygous Y genes respectively lost their Y- and X-coding sequence, while diploid are genes still having a copy on both chromosomes. For diploid genes, strata were defined using the Y copy position. Confidence intervals from 1000 bootstraps are shown. Sample sizes for each median are indicated in Table S10.

Apart from chromosome degeneration, gene expression on sex chromosomes is expected to evolve through several processes. First, as the X chromosome is more often transmitted to females, it is expected to accumulate dominant female-beneficial mutations that could lead to an increase in expression of the X copy (Bachtrog et al., 2011). Sex chromosomes are also expected to be enriched in sexual conflicts, in which case gene expression should increase depending on the sex in which they are beneficial (Vicoso & Charlesworth, 2006; Bachtrog et al., 2011). Finally, the lack of recombination and lowered sample size of the Y chromosome can lead to the accumulation of loss-of-function mutations (Bachtrog, 2013), leading to the progressive loss of Y-copy expression (Charlesworth & Charlesworth, 2000). We observed a feminization of stratum I in gills and II in brain, which had previously been described for stratum II (Leder et al., 2010; White et al., 2015), suggesting a role for female-beneficial mutations in the evolution of gene expression on the X chromosome. We did not observe feminization of the pseudo-autosomal region as previously reported by White et al (2015). This could be caused by the use of autosomal average expression level as the ancestral level of expression instead of the use of a closely related species to estimate each gene ancestral expression rate, which is more accurate.

To better understand the drivers of sex-biased gene expression of sex chromosomes, we compared expression of X and Y alleles in sex-biased genes with conserved copies on both chromosomes (excluding the still recombining PAR) to the allelic expression of unbiased genes (Fig. 6). In all tissues, we found a lowered expression of the Y allele of female-biased genes compared to unbiased genes (Wilcoxon rank-sum test p-value: 1×10^−4^ in brain, 8×10^−4^ in gills and 2×10^−2^ in liver) while X expression remained similar (all p-value > 0.7), which also resulted in lowered expression in males in the brain (p-value 1×10^−2^). This suggests that the degeneration of the Y chromosome coupled with the absence of dosage compensation in this species is the main driver of female-biased gene expression, as suggested by previous work (White et al., 2015). Note that we also found that in the liver overexpression in females also occurred in female-biased genes (p-value = 2×10^−2^).

**Figure 6:**
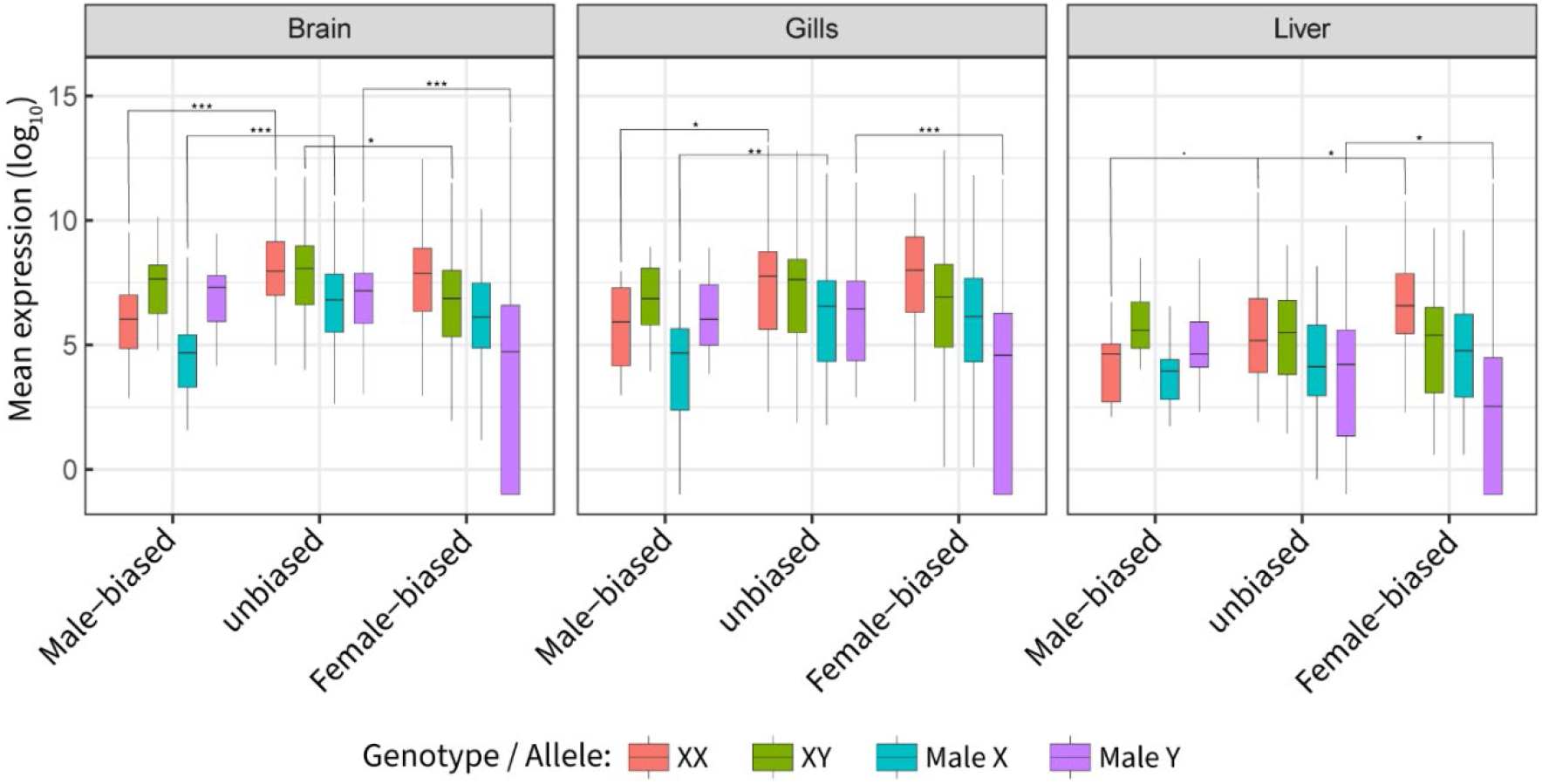
Genotype and allele-specific gene expression across male-biased, unbiased and female-biased genes. Male-biased and female-biased genes were defined as genes with log2 fold change in expression under −0.5 and above 0.5. We assessed significance using Wilcoxon rank-sum test comparing within genotype expression levels of male-biased and female-biased genes to unbiased genes. ^*^: p-value <0.05, ^**^ p-value <0.01, ^***^ p-value <0.001

Similarly, we observed that male-biased gene expression in females (6×10^−5^, 1×10^−2^ and 7×10^−2^) but not males (all p-value > 2×10^^−1^) was reduced compared to unbiased genes. We also observed reduced expression of the X allele in males in the brain and gills (1×10^−4^, 7×10^−3^), suggesting that this could be the result of a systematic down-regulation of some genes on the X-chromosome, which could be a signal of ongoing demasculinization, i.e. the loss of male-advantageous gene on the X chromosome (Gurbich & Bachtrog, 2008). Concordant with that hypothesis, genes identified as specific to the Y chromosome tended to have higher expression than genes specific to the X chromosome in males, suggesting that they are associated with male-beneficial functions.

## Conclusion

Our study characterized the sex-specific transcriptome of brain, gills and liver for wild three-spined sticklebacks during its reproductive period. We find low levels of differentiation between sexes in the brain compared to the level of dimorphism shown in behavioral and morphological studies. On the opposite, the gills exhibited pronounced sexual dimorphism that is usually not reported or accounted for in the literature, suggesting that the importance of sex as a cofactor in the study of gill physiology and function has been underestimated. The liver appeared to be strongly differentiated between the sexes, as expected for teleost fish because of its involvement in reproduction and metabolism. Sex chromosomes were a hotspot of intersex differentiation in all tissues, with ∼70% of genes being differentially expressed. This pattern seems to be caused both by an ongoing degeneration of the non-recombining region of this sex chromosome, coupled with the absence of dosage compensation mechanism, and a potential repression of male-advantageous mutations on the X chromosome, although further investigation of gene sequence evolution would be necessary.

## Acknowledgement

Authors are thankful to I. Caza-Allard E. Reni-Nolin for their help during fieldwork and tissue dissection. We thank S. Delaive and F-A Deschênes-Picard for a second year of sampling, although not used in this manuscript. We also thank S. Bernatchez, R. Bouchard, A. Perreault-Payette and C. Berger for occasional assistance and advice during fieldwork. We thank B. Bougas for her help during mRNA extraction and library construction, and Y. Dorant and E. Normandeau for their help during data processing. We also thank C. Venney for english proofing the manuscript. This project is part of the Ressources Aquatiques Québec research program. L. Bernatchez, who co-conceptualized and co-supervised this project as the PhD advisor of the first author, passed away before the end of it. We are extremely grateful for his involvement, support and motivation until the end.

## Data accessibility

Raw sequencing reads and unfiltered read counts are available though NCBI GEO, accession GSE269432. All analysis in the manuscript and related code are available at https://doi.org/10.5281/zenodo.11477976. The list of all sex-biased genes for each tissue is avaialable as supplementary table 11.

