## Supplementary material for "Sex-biased gene expression across tissues reveals unexpected differentiation in the gills of the threespine stickleback": Figure S1

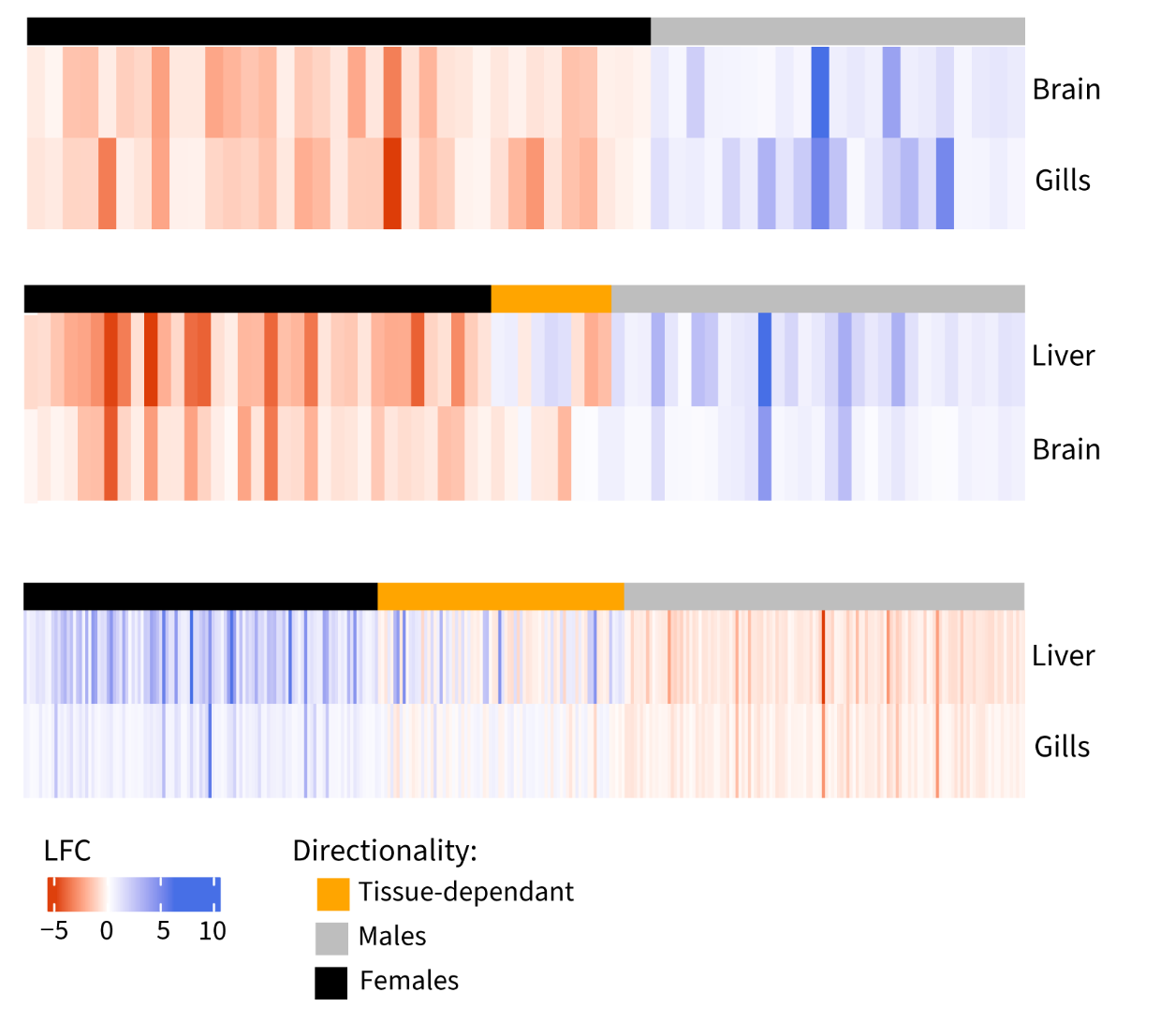

Supplementary Figure 1: Heatmap of log-fold change in gene expression between sexes for genes significantly differentially expressed in pairs of tissue. Bar on top shows concordance in the direction of expression bias. From top to bottom: Gene significant in both brain and gills, liver and brain, and liver and gills.
